# Integrative and Conjugative Elements (ICE) and Associated Cargo Genes within and across Hundreds of Bacterial Genera

**DOI:** 10.1101/2020.04.07.030320

**Authors:** James H Kaufman, Ignacio Terrizzano, Gowri Nayar, Ed Seabolt, Akshay Agarwal, Ilya B Slizovskiy, Noelle Noyes

## Abstract

Horizontal gene transfer mediated by integrative and conjugative elements (ICE) is considered an important evolutionary mechanism of bacteria. It allows organisms to quickly evolve new phenotypic properties including antimicrobial resistance (AMR) and virulence. The rate of ICE-mediated cargo gene exchange has not yet been comprehensively studied within and between bacterial taxa. In this paper we report a big data analysis of ICE and associated cargo genes across over 200,000 bacterial genomes representing 1,345 genera. Our results reveal that half of bacterial genomes contain one or more known ICE features (“ICE genomes”), and that the associated genetic cargo may play an important role in the spread of AMR genes within and between bacterial genera. We identify 43 AMR genes that appear only in ICE genomes and never in non-ICE genomes. A further set of 95 AMR genes are found >5x more often in ICE versus non-ICE genomes. In contrast, only 29 AMR genes are observed more frequently (at least 5:1) in non-ICE genomes compared to ICE genomes. Analysis of NCBI antibiotic susceptibility assay data reveals that ICE genomes are also over-represented amongst phenotypically resistant isolates, suggesting that ICE processes are critical for both genotypic and phenotypic AMR. These results, as well as the underlying big data resource, are important foundational tools for understanding bacterial evolution, particularly in relation to important bacterial phenotypes such as AMR.

## Background

Several mechanisms of horizontal gene transfer (HGT) allow bacteria to exchange genetic code. One of these mechanisms, termed conjugation, occurs when bacterial cells form direct physical contacts that allow for passage of genetic material from one bacterium to another. The machinery required to form these contacts and initiate genetic exchange is often contained within integrative and conjugative elements (ICE) [1, 2, 3, 4, 5, 6, 7]. ICE are modular mobile genetic elements that integrate into host genomes; are propagated via cellular replication; and can be induced to excise from the host genome in order to initiate the process of conjugation. The conditions that induce excision and conjugation are not fully elucidated, but DNA damage and subsequent SOS response seem to be an important trigger [8, 9].

Genes exchanged between bacteria during ICE-mediated transfer include functional domains associated with ICE machinery (e.g. excisionases, integrases, conjugative transport proteins, etc...) as well as intervening ‘accessory’ sequences that encode a variety of cargo genes [6]. By pairing ICE machinery with an array of diverse cargo genes, bacterial communities can significantly expand their genetic repertoire, including between bacteria of diverse taxonomy [6, 10, 11, 12]. Functions commonly associated with ICE cargo include antimicrobial resistance (AMR) and virulence [3, 5, 6, 10], both of which represent risks to human and animal health if transferred into pathogens. Therefore, understanding the microbial ecology of ICE and cargo genes (i.e., their distribution and behavior across bacterial taxa) is important in assessing the human health risk posed by various bacterial communities. For example, how often do different bacterial taxa carry ICE and AMR genes; what resistance phenotypes are commonly associated with ICE machinery; how often do different commensal bacterial taxa use ICE to exchange various cargo genes with pathogens; and what conditions foster ICE-mediated exchange of specific cargo genes between pathogens and non-pathogens? These questions are fundamental to understanding how bacterial communities respond to external stimuli, and how these responses increase the overall risk posed by microbial communities of varying composition [13, 14]. Understanding these dynamics, in turn, allows us to better understand how human practices may increase “risky” microbial behaviors (such as ICE-mediated exchange), and thus predispose to higher-risk microbial communities. For example, we can start to predict how antimicrobial use practices impact the likelihood of pathogens obtaining AMR genes from the commensal microbiome via ICE transfer.

These microbial ecological questions are becoming increasingly tractable as more and more whole genome sequence (WGS) data are generated. As an example, the analysis of HGT-associated genes from just 336 genomes across 16 phyla was sufficient to significantly improve bacterial phylogenies as compared to those obtained from conserved marker genes [15]. An analysis of 1,000 genomes demonstrated that ICE machinery is ubiquitous across diverse prokaryotes, and likely one of the most common mechanisms of bacterial evolution [11]. Today, public datasets contain orders of magnitude more WGS data. However, despite the importance of HGT in bacterial evolution and pathogenicity, there has not yet been a comprehensive, systematic survey of the frequency of ICE and cargo protein sequences within or between bacterial genera. The objective of this work was to describe intra- and inter-genus ICE-cargo dynamics using the comprehensive set of WGS data and ICE sequences currently available.

Using both sequence- and annotation-based queries of a relational database developed from the National Center for Biotechnology Information (NCBI), we report a large-scale computational analysis of ICE and associated cargo proteins across 186,887 non-redundant bacterial genomes representing over 1,300 genera. Using 36 ICE features (Table 4) representing different families of ICE (i.e., specific proteins or protein families from UniProtKB and the ICE literature), we identify 95,781 genomes that contain at least one ICE feature. We term these “ICE genomes”, and note that they represent 631 of the 1,345 analyzed genera (47%). In a detailed analysis of potential ICE-mediated exchange within and between genera based on exact-match sequence similarity, we find that the ICE genomes contain 28,042 distinct ICE proteins and 11,276,651 corresponding cargo proteins (out of 51,362,178 total unique protein sequences in the source database). The full set of cargo genes map to 20,550 distinct gene names (excluding ‘putative protein’ or ‘hypothetical protein’), with a wide range of ICE-mediated transfer frequencies within and between genera.

To gain insight into ICE cargo genes, we perform a statistical comparison of all genes annotated with names associated with AMR. By comparing the frequency with which these genes appear in ICE genomes versus non-ICE genomes, we find that out of 286 AMR gene names, 220 are found more often in ICE genomes and 63 are found more often in non-ICE genomes (and 3 with equal probability). Furthermore, we find that rare or less frequently observed AMR genes are more likely to be associated with ICE genomes, while common or abundant genes are less likely to be associated with ICE genomes. In an independent analysis of phenotypic antibiotic susceptibility data contained in NCBI BioSample data, we evaluate all public assay data for ICE and non-ICE genomes. Considering all antibiotic drug compounds with more than 60 phenotypic resistant measurements, the data show that resistance occurs in ICE genomes with probability >80% *regardless of compound*. By comparison, in a random process, the probability would be expected to be closer to 50% based on the prevalence of ICE genomes.

These results advance our understanding of the complex microbial ecological dynamics arising from ICE-mediated exchange, and represent a significant “big data” resource for scientists working on microbiome research and pan-microbial evolution. As such, our results have wide-reaching impact, spanning from theoretical underpinnings of microbial behavior to infectious disease and food safety.

## Results

Determination of ICE genomes, ICE proteins, and those proteins with the greatest supporting evidence as possible cargo proteins is described in detail in *Methods*. To test the identification of ICE and cargo proteins we use the annotated locus to ensure that the ICE protein features occur within regions of putative cargo (and not surrounded by other chromosomal genes). Figure 1 shows that the vast majority of putative cargo proteins are adjacent within contigs, and that the contigs containing the ICE features themselves are more likely to be proximate to putative cargo proteins (black regions) vs non-cargo proteins (grey regions). We note that each genome in the figure is actually a linear representation of the contigs assembled by SPAdes, in the order in which they were annotated by Prokka. Due to inherent genome characteristics and the nature of short-read sequence data, SPAdes (and other *de novo* genomic assemblers) can not always establish the assembly order for all contigs. Therefore, when contigs containing no cargo proteins are interspersed with cargo-containing contigs, we can not definitively determine whether this is due to fragmented assemblies or biology. Such interspersion shows up as noise within the (black) regions of likely cargo proteins; this is best observed by viewing the heatmap in full screen. We emphasize that the noise is not evidence for or against chromosomal rearrangement after ICE exchange, but rather could be artifact of the contig-centric annotation process.

**Figure 1.**
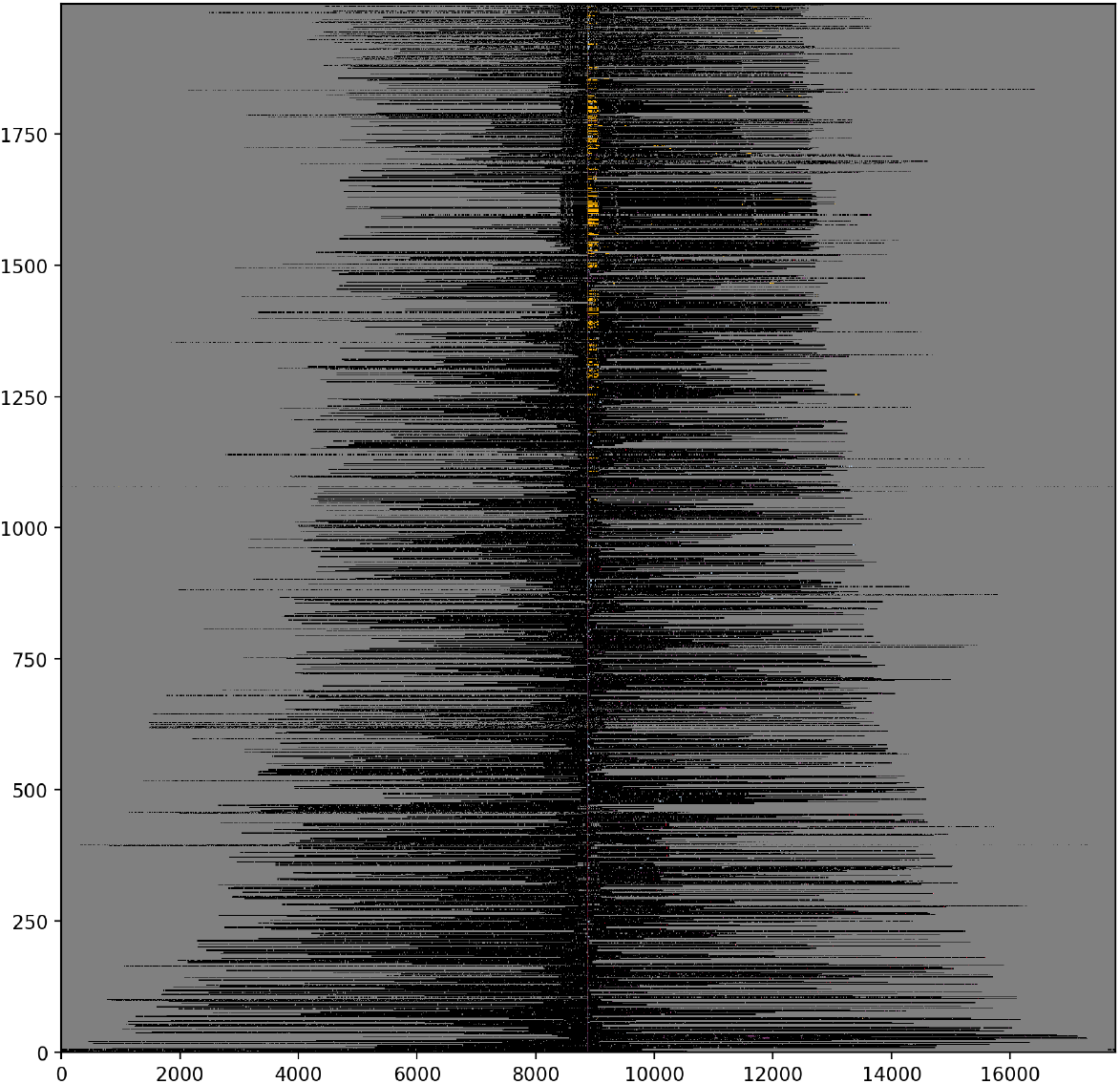
Heatmap with relative positions of ICE features. Heat map showing the relative position of ICE features and putative cargo proteins within contigs for the 2,000 genomes with the greatest number of cargo proteins. ICE features are represented as color pixels based on the colors shown in Table 4. Cargo proteins are shown in black and other chromosomal DNA in grey. Each genome is bit shifted to the left until the first ICE feature is centered in the figure. Most genomes contained more than one ICE protein (see text).

Figure 1 is intended to test if the observed ICE features (Table 4) co-occur proximate to regions of cargo proteins (shown in black). The figure shows the relative position of cargo and ICE proteins within assembled contigs. Each row in the figure represents a genome, with ICE proteins shown in color based on Table 4, cargo proteins in black, and other sections of the genome in grey. Each genome’s data are rotated (bit shifted) left so as to center the first ICE protein, which appears as a colored, vertically-aligned pixel. For genomes containing more than one ICE protein, additional ICE proteins appear as individual scattered colored pixels to the right of the central line. In the supplement we provide a similar figure for each ICE protein class listed in table 4.

From Figure 1, it is also evident that individual genomes may contain more than one ICE protein, and that these proteins may in turn represent more than one ICE feature (Table 4). Figure 2 plots the number of ICE features per genome (a), and the number of ICE proteins per genome (b) as a function of the number of genomes on a logarithmic scale. Note that the genome order in (a) and (b) is different and selected to sort the feature counts from greatest to least in each case.

**Figure 2.**
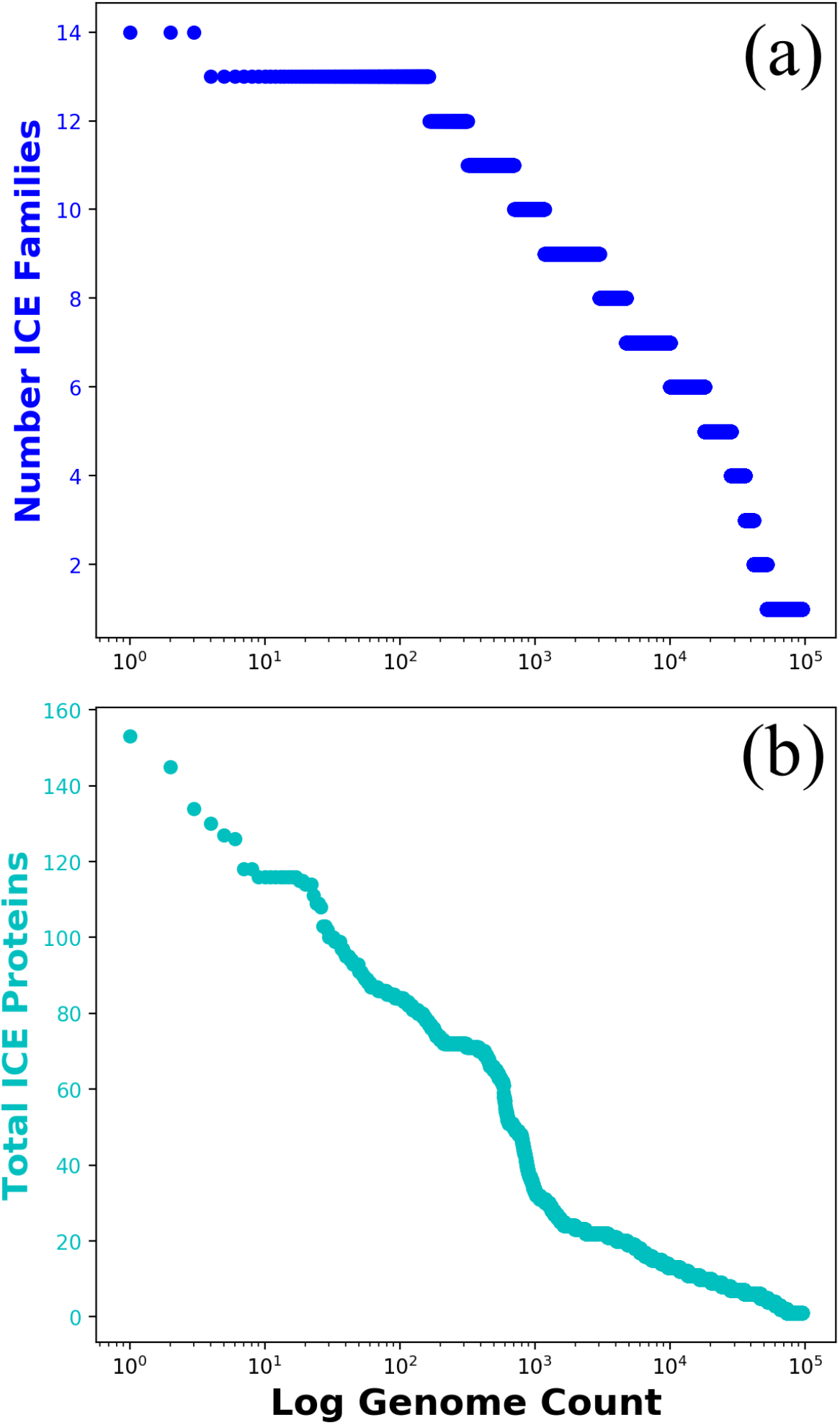
ICE Features per Genome. Individual genomes typically contain more than one ICE feature (Table 4, and often contain more than one protein per feature. Figure 2 (a) shows the number of ICE features per genome, and Figure 2 (b) shows the number of ICE proteins per genome for all 106,433 genomes containing at least one ICE feature. For both panels, the genome order is sorted by number of features (largest to smallest).

A priori, one might reasonably expect the number of observed ICE genomes to scale with the number of genomes available for each genus. However, genus representation in NCBI is not uniform across genera, leading to bias in available genomes per genus. To correct for this imbalance, we computed the ICE genome frequency by normalizing the number of observed ICE genomes to the number of genomes per genus. Table 1 lists all genera with over 100 representative genomes in OMXWare, ordered by the proportion of ICE genomes in the genus. The genera with the largest fraction of ICE genomes are *not* the genera with the most genomes in NCBI. For example, although *Salmonella* has by far the greatest number of high quality genomes (N=39,574), it ranks fifth in terms of the proportion of genomes that contain an ICE protein. The top 30 genera listed in Table 1 all have an ICE genome frequency greater than 20%, with genomes from the genus *Legionella* containing ICE proteins over 99% of the time. This high percentage may be due to sampling bias in the available NCBI WGS datasets (for example, the *Legionella pneumophila* WGS accessions appear to have been collected from a single site), or it may represent the propensity for ICE-mediated processes to occur within individual genera.

**Table 1.** Frequency of ICE genomes, by genus. The table lists all genera with over 100 representative genomes ordered by the frequency of ICE genomes in the genus. The green line separates ICE frequency above or below 50%.

| Genus | ICE Genomes | Total Genomes | ICE Genome Frequency |
| --- | --- | --- | --- |
| Legionella | 1672 | 1686 | 0.99 |
| Shigella | 5423 | 5541 | 0.98 |
| Klebsiella | 4682 | 5304 | 0.88 |
| Elizabethkingia | 102 | 119 | 0.86 |
| Escherichia | 8140 | 9957 | 0.82 |
| Stenotrophomonas | 441 | 563 | 0.78 |
| Enterobacter | 894 | 1210 | 0.74 |
| Vibrio | 2902 | 4017 | 0.72 |
| Acinetobacter | 2621 | 3770 | 0.70 |
| Pseudomonas | 3222 | 4750 | 0.68 |
| Enterococcus | 1003 | 1516 | 0.66 |
| Citrobacter | 131 | 203 | 0.65 |
| Salmonella | 24123 | 38808 | 0.62 |
| Clostridioides | 1329 | 2183 | 0.61 |
| Streptococcus | 8244 | 13766 | 0.60 |
| Xanthomonas | 201 | 357 | 0.56 |
| Staphylococcus | 18034 | 32661 | 0.55 |
| Rhizobium | 110 | 202 | 0.54 |
| Yersinia | 215 | 437 | 0.49 |
| Lactococcus | 57 | 117 | 0.49 |
| Serratia | 230 | 619 | 0.37 |
| Sinorhizobium | 44 | 121 | 0.36 |
| Bifidobacterium | 137 | 403 | 0.34 |
| Moraxella | 65 | 192 | 0.34 |
| Bacillus | 471 | 1471 | 0.32 |
| Campylobacter | 5340 | 19501 | 0.27 |
| Aeromonas | 80 | 312 | 0.26 |
| Brucella | 230 | 970 | 0.24 |
| Mesorhizobium | 89 | 385 | 0.23 |
| Helicobacter | 118 | 529 | 0.22 |
| Streptomyces | 74 | 333 | 0.22 |
| Corynebacterium | 133 | 639 | 0.21 |
| Neisseria | 153 | 781 | 0.20 |
| Burkholderia | 341 | 2053 | 0.17 |
| Haemophilus | 65 | 403 | 0.16 |
| Lactobacillus | 144 | 962 | 0.15 |
| Listeria | 848 | 7716 | 0.11 |
| Clostridium | 40 | 454 | 0.09 |
| Mycobacterium | 1120 | 13129 | 0.09 |
| Cutibacterium | 10 | 118 | 0.08 |
| Bordetella | 60 | 733 | 0.08 |
| Chlamydia | 33 | 496 | 0.07 |
| Bartonella | 3 | 124 | 0.02 |
| Mycoplasma | 2 | 251 | 0.01 |
| Francisella | 0 | 120 | 0.00 |

In order to minimize false positive identification of ICE and potential cargo proteins, we applied a strict rule that ICE proteins must be seen with identical amino acid sequence in two or more genomes, and that cargo proteins must be seen in two or more ICE genomes with exact amino acid sequence identity. The strict sequence identity rule likely decreases the sensitivity to detect ICE-mediated transfer events, as genomes continue to evolve after HGT events, including possible chromosomal rearrangement [16]. However, if ICE-mediated HGT occurs frequently enough, and if our reference database is large enough that we can rely on the “law of large numbers” [17], then the set of all genomes should contain some *pairs of genomes* that have not yet evolved so far as to obscure the cargo proteins. Figure 1 is a test of this hypothesis, and provides evidence that cargo proteins identified by the strict selection process exist in genomic proximity to each other and to the ICE proteins that likely transferred them. This rule is also required to visualize the transfer of ICE and cargo proteins within and between genera, as in Figure 3. In this force-directed graph visualization, each node is a genus and edges represent inter-genus transfer. Edges can be filtered based on various criteria, including type of ICE involved in the transfer; number of ICE proteins; distinct cargo proteins (by sequence); and the number of AMR cargo proteins by sequence. By definition, edges in Figure 3 require sequence identity at each node, i.e., genus. Edge weights represent either (1) the number of ICE proteins shared between the genera, (2) the number of distinct proteins shared between the genera, or (3) the number of AMR proteins shared between the genera – depending on filtering criteria used. The size of nodes in Figure 3 represents the fraction of genomes within that genus identified as ICE genomes. Full data are available in tabular form and the visualization is available in the supplement.

**Figure 3.**
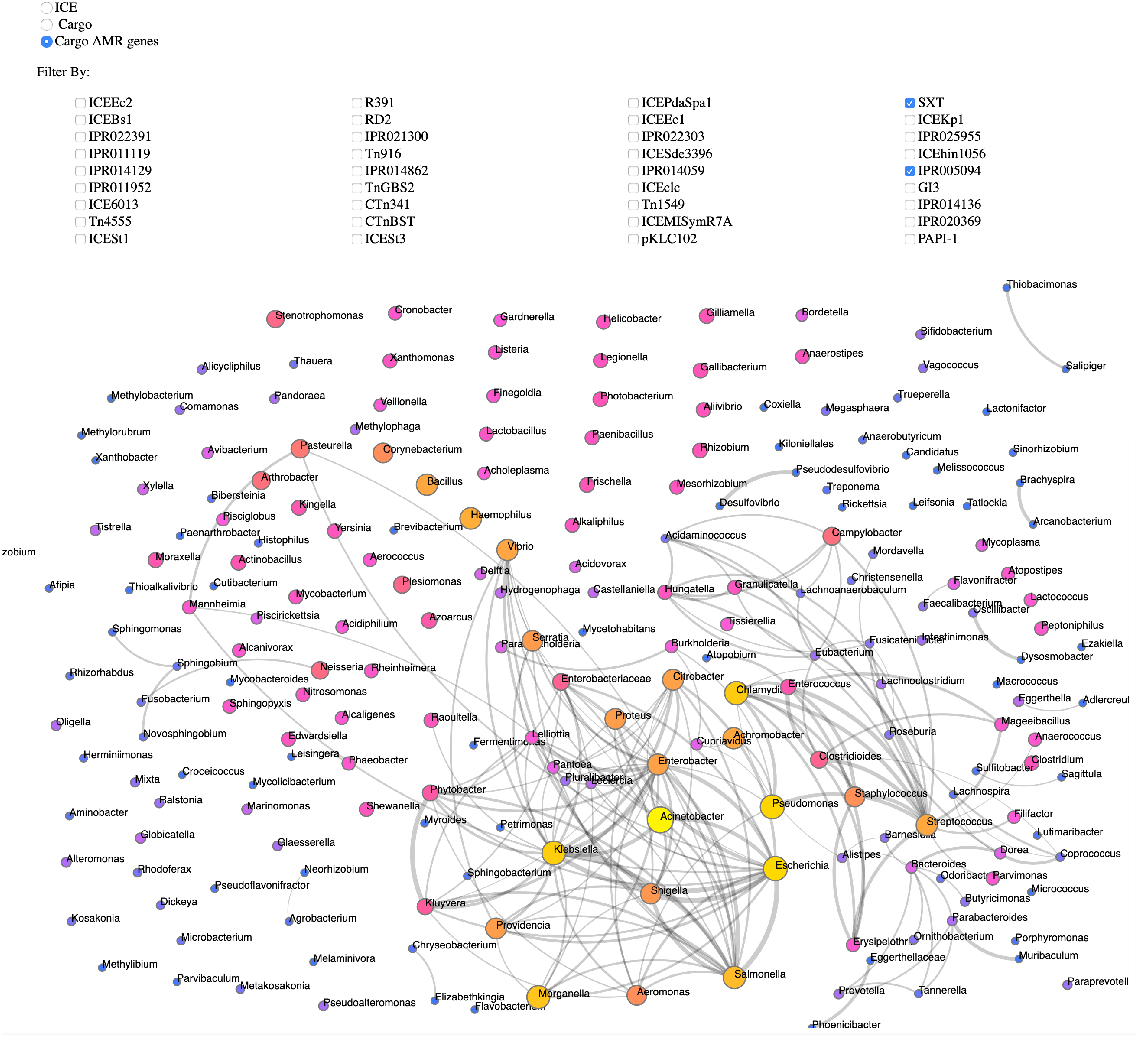
Force-Directed Graph. Force-directed graph showing the frequency of ICE and cargo proteins shared within and between genera. *See: large format animation in supplement*

A *subset* of the observed cargo protein names are associated with a set of confirmed antimicrobial resistance (AMR) protein names. We identified this subset by selecting *only* those names that Prokka assigned to sequences mapping to a name defined in MEGARes v1.0 [18]. The entity relations in our database ensure a 1::1 mapping between gene and protein names and their respective sequences. Of the 3,824 distinct sequences contained in MEGARes, Prokka identified 3,674 of them as valid sequences coding for protein. These 3,674 distinct proteins were assigned 286 distinct names, excluding ‘putative protein’ and ‘hypothetical protein’. While this highly curated set is certainly not a comprehensive list of all proteins contributing to AMR, it is a useful initial set to estimate the fraction of AMR proteins within the larger set of ICE and cargo proteins.

We analyzed this set of AMR proteins in an attempt to estimate or bound the number of possible false classifications of cargo proteins that are specifically related to ICE-mediated HGT. This was necessary because some genomic features are shared between ICE and plasmids, an indeed some of the InterProScan codes that define ICE features may also contain machinery required for the formation or integration of plasmids. Accordingly, Figure 4 shows a 2-d histogram of AMR protein sequences (by AMR name). Along the x-axis, the histogram represents the frequency with which AMR protein sequences are found on ICE vs non-ICE genomes. Along the y-axis, the histogram represents the *fraction* of the protein sequences (by AMR name) that have been observed on plasmids. Note the full scale on the y-axis. The maximal frequency of sequence observation on a plasmid is 10%, and this provides an upper bound to possible false positive identification as ICE-related cargo. In fact, observation of a sequence required for plasmid formation on a plasmid does not rule out observation of the same sequence on ICE, and vice versa.

**Figure 4.**
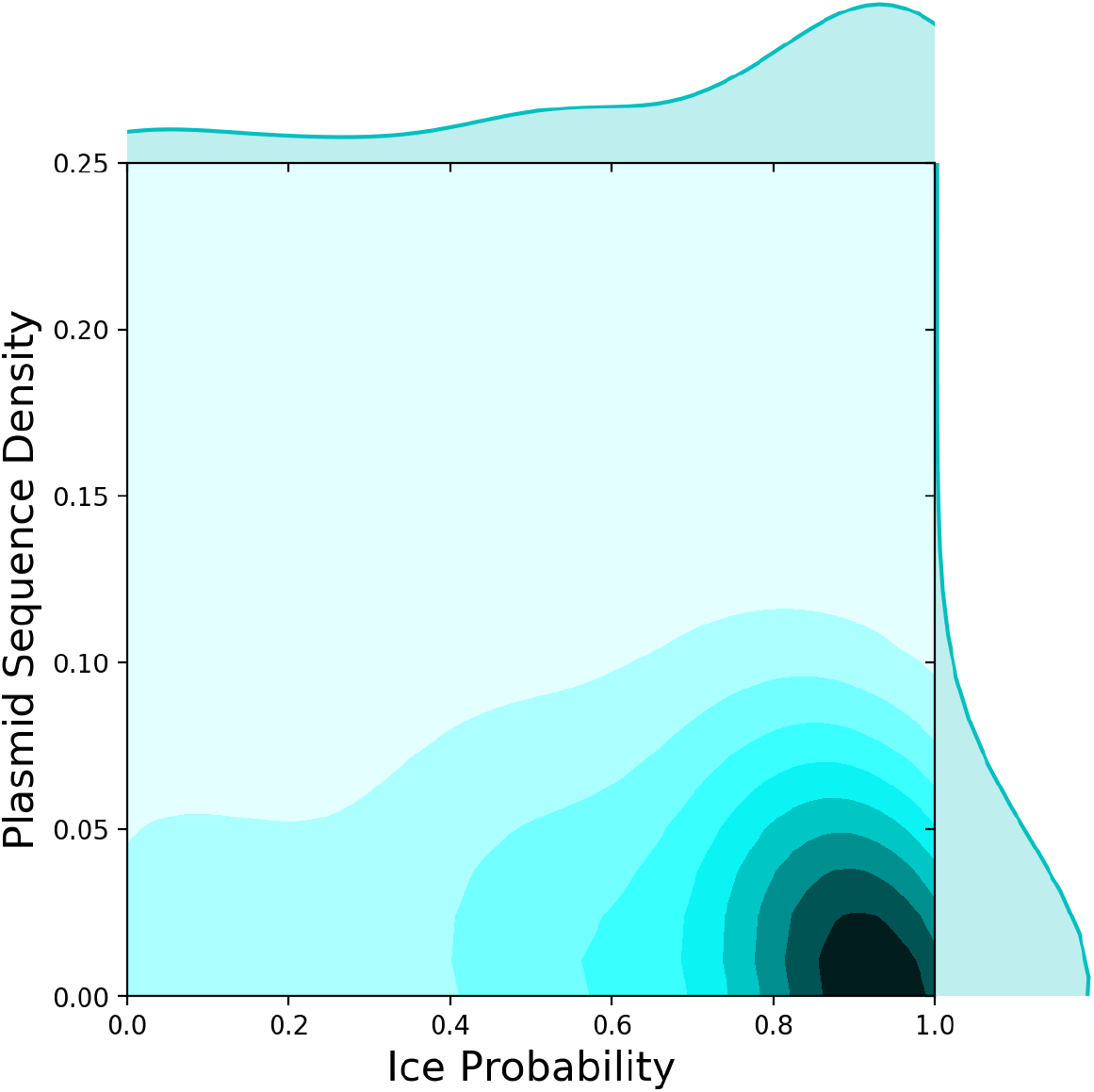
Probability AMR Genes Found in ICE Genomes. 2-d Histogram showing the probability that AMR proteins (by name) are found in ICE Genomes (x-axis) and the fraction of AMR protein sequences (by name) observed on plasmids (y-axis). Independent of ICE probability, 5-10% of AMR sequences have been observed on plasmids. The majority of AMR proteins are found on ICE genomes.

To analyze the distribution of AMR proteins across genomes, we calculated the probability that each AMR gene name was identified in ICE versus non-ICE genomes, (Table 2. From the complete list of 286 AMR protein names (see *supplement*), 220 are found more often in ICE genomes, 3 are found in ICE and non-ICE genomes with equal probability, and 63 are found more often in non-ICE genomes. Given that the ratio of ICE:non-ICE genomes in our database is 51%, this suggests that AMR genes are disproportionately represented within ICE genomes.

**Table 2.** Probability of observing AMR names in ICE vs non-ICE genomes. Only those AMR names that appear in more than 100 genomes are considered. The table shows those AMR names that occur with ICE genome probability ≥ 0.98 (above line) and ≤ 0.15 (below line). The full data is available in the online supplement.

| AMR Protein Name | Total Genome Count | ICE Genome Count | non-ICE Genome Count | ICE Genome Prob |
| --- | --- | --- | --- | --- |
| Rob DNA-binding transcriptional activator | 4559 | 4559 | 0 | 1.00 |
| Transposon Tn10 TetD protein | 4559 | 4559 | 0 | 1.00 |
| Tetracycline resistance ribosomal protection protein Tet(M) | 304 | 304 | 0 | 1.00 |
| Outer membrane protein YedS | 288 | 288 | 0 | 1.00 |
| Regulator of RpoS | 288 | 288 | 0 | 1.00 |
| Tetracycline resistance protein TetM | 137 | 137 | 0 | 1.00 |
| Beta-lactamase Toho-1 | 319 | 318 | 1 | 1.00 |
| AcrAD-TolC multidrug efflux transport system - permease subunit | 1584 | 1576 | 8 | 0.99 |
| Multidrug efflux pump subunit AcrB | 1584 | 1576 | 8 | 0.99 |
| DNA topoisomerase subunit A | 1091 | 1085 | 6 | 0.99 |
| MATE family multidrug efflux pump protein | 1607 | 1596 | 11 | 0.99 |
| Carbapenem-hydrolyzing beta-lactamase KPC | 1914 | 1899 | 15 | 0.99 |
| Beta-lactamase OXA-1 | 1188 | 1175 | 13 | 0.99 |
| Inner membrane protein HsrA | 350 | 346 | 4 | 0.99 |
| Putative transport protein YdhC | 342 | 338 | 4 | 0.99 |
| Aclacinomycin methylesterase RdmC | 419 | 414 | 5 | 0.99 |
| Beta-lactamase OXA-2 | 199 | 196 | 3 | 0.98 |
| Chloramphenicol efflux MFS transporter CmlA1 | 795 | 783 | 12 | 0.98 |
| Beta-lactamase OXA-10 | 521 | 513 | 8 | 0.98 |
| 16S rRNA (guanine(1405)-N(7))-methyltransferase | 994 | 978 | 16 | 0.98 |
| Ribosomal RNA large subunit methyltransferase H | 4087 | 3989 | 98 | 0.98 |
| Multidrug resistance operon repressor | 486 | 67 | 419 | 0.14 |
| Outer membrane protein OprM | 478 | 65 | 413 | 0.14 |
| Methicillin-resistance regulatory protein MecR1 | 8344 | 1007 | 7337 | 0.12 |
| Phosphoethanolamine-lipid A transferase MCR-1.1 | 8344 | 1007 | 7337 | 0.12 |
| HTH-type transcriptional repressor BepR | 467 | 55 | 412 | 0.12 |
| Bifunctional polymyxin resistance protein ArnA | 507 | 53 | 454 | 0.10 |
| Methicillin resistance regulatory protein MecI | 6583 | 395 | 6188 | 0.06 |
| Metallothiol transferase FosB | 6584 | 395 | 6189 | 0.06 |
| Multidrug efflux transporter MdtL | 419 | 9 | 410 | 0.02 |
| Multidrug efflux pump subunit AcrA | 420 | 7 | 413 | 0.02 |
| HTH-type transcriptional regulator SyrM 1 | 414 | 3 | 411 | 0.01 |
| Multidrug efflux RND transporter permease subunit OqxB3 | 358 | 2 | 356 | 0.01 |
| Aminoglycoside 2'-N-acetyltransferase | 9723 | 5 | 9718 | 0.00 |
| DNA-binding response regulator MtrA | 9854 | 2 | 9852 | 0.00 |
| Putative acetyltransferase | 5232 | 1 | 5231 | 0.00 |
| Quinolone resistance protein NorB | 213 | 0 | 213 | 0.00 |

The analysis above does not distinguish between different ICE features, and treats AMR proteins as independent features. However, the data in Figure 1 demonstrate that many of the ICE features defined in Table 4 often co-occur in the same genome, as do some of the AMR proteins. To gain insight into these correlations, and to identify groups of AMR proteins associated with different ICE families, we show in Figure 5 a co-occurrence matrix across all ICE features for the 43 AMR proteins with an ICE:non-ICE genome frequency of >5x. This figure demonstrates that some ICE features co-occur within genomes frequently with both other ICE features as well as multiple AMR protein names. For example, IPR005094 appears to co-occur with the widest diversity of AMR protein names. Many co-occurrence patterns in Figure 5 reflect known biological associations. For example, the Tn916 ICE feature co-occurs most frequently with tetracycline ribosomal protection protein TetM, a genomic association discovered over three decades ago [19]. While TetM seems to co-occur with a few select ICE features (such as Tn916), other AMR protein names seem to co-occur with many ICE features. For example, many of the AMR names associated with extended-spectrum beta-lactam and carbapenem resistance co-occur with the majority of evaluated ICE features (e.g., beta-lactamases Toho-1, OXA-1, OXa-2, OXA-10, SHV-2, and KPC), which may provide partial explanation for the observed rapid expansion of these important AMR proteins within Enterobacteriaceae [20]. Similarly, the recently widely-publicized mcr-1 protein seems to co-occur with multiple ICE features, which both strengthens and expands upon recent findings that this AMR gene has been mobilized on numerous plasmid types [21]. Co-occurrence data such as those provided in Figure 5 may represent a new and sustainable (i.e., easily updated) source of information regarding the potential for new and emerging AMR genes to expand within and across bacterial populations. This information, in turn, could help to prioritize and focus public health and human clinical decisionmaking regarding AMR.

**Figure 5.**
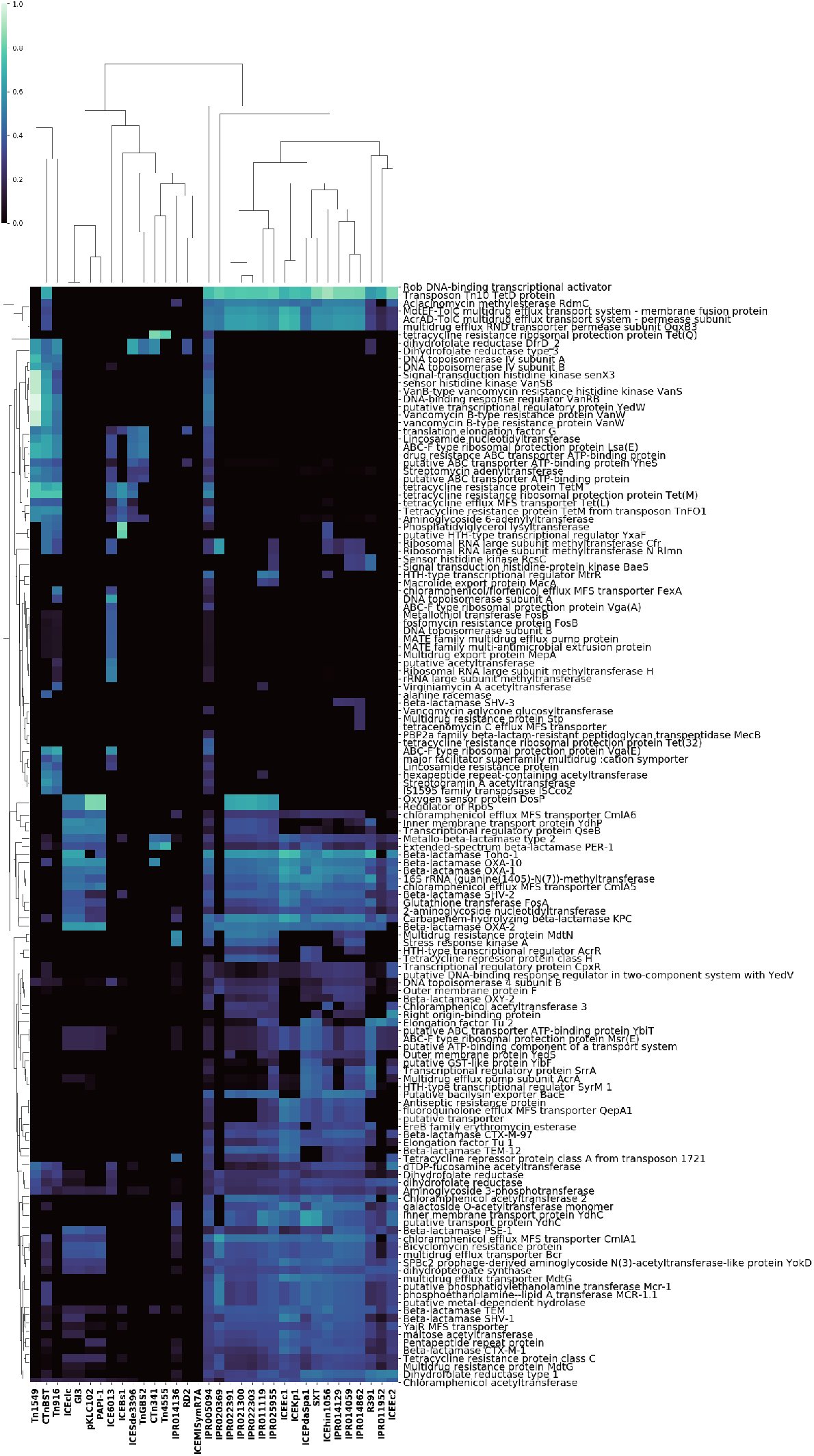
Co-occurrence of 138 AMR proteins (by name) with ICE features. Co-occurrence of AMR proteins (by name) with ICE features, for the 138 AMR proteins that are observed in ICE genomes with a frequency greater than 5x the frequency of observation in non-ICE genomes. Of these proteins, 43 are *only* observed in ICE genomes.

Conversely, in Figure 6 we display a co-occurrence matrix for all 29 AMR proteins observed in non-ICE genomes with a frequency greater than 5x the frequency of observation in ICE genomes. Of note is the observation that no beta-lactam AMR protein names occur in this list of 29 AMR names; this contrasts starkly to the preponderance of beta-lactam-associated AMR names in Figure 5, again suggesting that beta-lactam resistance is tightly coupled with ICE machinery, and that ICE-mediated exchange is the primary evolutionary driver of beta-lactam resistance. By comparison, several mechanisms of multi-drug resistance (MDR) are contained within the list of 29 AMR proteins observed more frequently in non-ICE versus ICE genomes, i.e., AcrB, AcrE, OqxB7, mdtA, mdtE, mdtH, and MexB. These mechanisms of MDR tend to be multi-purpose, i.e., the proteins confer multiple functional benefits to bacteria, in addition to AMR. Together, the results of Figures 5 and 6 suggest that proteins with more specific AMR functions tend to be disproportionately represented amongst ICE genomes, while more generalist proteins tend to be disproportionately represented within non-ICE genomes. One hypothesis for this observation is that the fitness cost-benefit dynamics differ for generalist versus specialist genes, such that specialized genes are more likely to transiently yet rapidly spread within bacterial populations via the so-called ‘accessory genome’ (which includes ICE-mediated exchange), whereas generalist genes are more likely to be maintained permanently within bacterial genomes, and thus are less likely to be identified as ICE-associated cargo.

**Figure 6.**
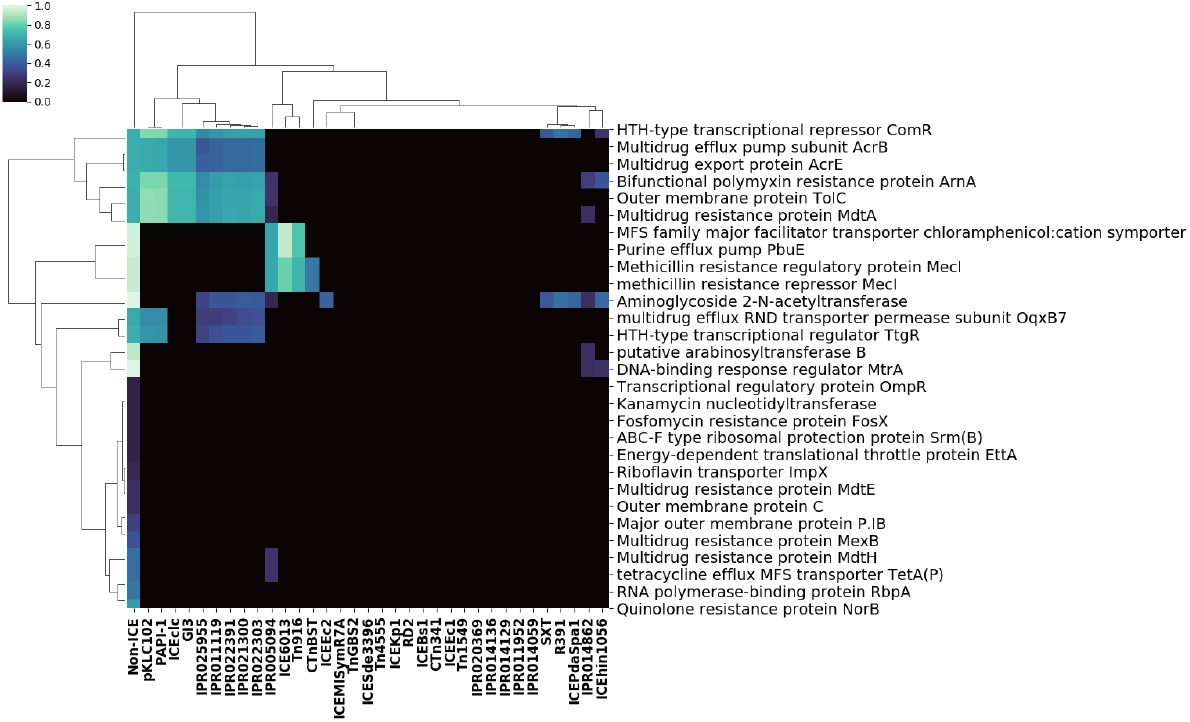
Co-occurrence of 29 AMR proteins (by name) with ICE features. Co-occurrence of AMR proteins (by name) with ICE features for the 29 AMR proteins observed in non-ICE genomes with a frequency at least 5x the frequency of observation in ICE genomes

Given our hypothesis that ICE-mediated spread of specialized AMR genes may be promoted by more specific evolutionary pressure such as antimicrobial drug exposures, we hypothesized that this signature of selective pressure may also manifest in the phenotypic properties of ICE versus non-ICE genomes. To evaluate this, we queried the NCBI BioSample assay metadata in our relational database (described in *Methods*), to identify isolates that had been tested for phenotypic antibiotic susceptibility (AST) to known antibiotic compounds. For the 186,887 highest quality genomes, the NCBI assay metadata contained 15,286 phenotypically-confirmed resistant genome-compound tests, representing 13,076 tests for ICE genomes and 2,210 tests for non-ICE genomes. Altogether, 1,242 genomes were used in these tests, of which 1,023 were ICE genomes and 219 were non-ICE genomes. For each antibiotic compound listed, we computed the number of phenotypically resistant isolates with ICE-vs non-ICE-containing genomes.

In Table 3 we list the probability of observing phenotypic resistance from ICE genomes, for all antibiotic compounds with more than 100 assays in the NCBI BioSample data. The full list is available in the supplement. The data reveal that phenotypic resistance occurs in ICE genomes with probability >80%, regardless of compound. As with the disproportionate representation of AMR genes within ICE genomes, the phenotypic AMR data again suggest that microbial AMR dynamics are driven largely by ICE-mediated processes. However, it is also important to note that NCBI phenotypic assay data is likely biased due to the motivations for clinicians and researchers to submit isolates for phenotypic testing. Therefore, to test for SRA sampling bias with respect to these compounds, we also measured the phenotypically-resistant fraction expected for *randomly selected* genomes, based on the number of genomes tested per compound in table 4 and the actual number of ICE and non-ICE genomes across the entire database. This null hypothesis was tested by running 100 bootstrapped trials for each compound. The results are listed in table 4. The average ICE probability for all compounds listed in table 4, weighted by total genomes tested per compound, is 0.85 ± .05 *independent of antibiotic compound*. In a random process, the probability would be expected to be near 51% given the fraction of all genomes with ICE features.

**Table 3.** The role of ICE in phenotypic resistance for specific antibiotic compounds. The table lists the probability of observing resistance in ICE genomes. Only compounds with 100 or more non-redundant resistant genome measurements are listed. The table shows that phenotypic AMR occurs in ICE genomes with probability >80% regardless of compound. The table also shows the number of expected resistant ICE genomes based on a bootstrapped random selection process with 100 trials (null hypothesis), using the number of assays and the actual fraction of genomes with ICE features (~51%).

| Antibiotic Compound | Total Resistant Genomes | ICE Resistant Genomes | Random Process | ICE Prob |
| --- | --- | --- | --- | --- |
| doripenem | 215 | 207 | $86 \pm 7$ | 0.96 |
| cefepime | 272 | 259 | $108 \pm 8$ | 0.95 |
| ampicillin-sulbactam | 433 | 402 | $173 \pm 10$ | 0.93 |
| imipenem | 429 | 396 | $171 \pm 11$ | 0.92 |
| piperacillin-tazobactam | 280 | 258 | $112 \pm 9$ | 0.92 |
| meropenem | 402 | 367 | $161 \pm 11$ | 0.91 |
| trimethoprim-sulfamethoxazole | 868 | 775 | $348 \pm 14$ | 0.89 |
| ertapenem | 222 | 197 | $89 \pm 8$ | 0.89 |
| levofloxacin | 741 | 644 | $297 \pm 14$ | 0.87 |
| gentamicin | 813 | 706 | $325 \pm 13$ | 0.87 |
| ciprofloxacin | 915 | 794 | $367 \pm 14$ | 0.87 |
| amoxicillin-clavulanic acid | 277 | 240 | $111 \pm 8$ | 0.87 |
| ceftriaxone | 984 | 849 | $394 \pm 16$ | 0.86 |
| tetracycline | 700 | 603 | $281 \pm 12$ | 0.86 |
| ceftazidime | 852 | 731 | $342 \pm 14$ | 0.86 |
| cefotaxime | 889 | 758 | $355 \pm 15$ | 0.85 |
| tobramycin | 674 | 574 | $270 \pm 12$ | 0.85 |
| ampicillin | 1042 | 852 | $418 \pm 16$ | 0.82 |
| amikacin | 383 | 313 | $154 \pm 10$ | 0.82 |
| aztreonam | 989 | 805 | $397 \pm 15$ | 0.81 |
| cefazolin | 1002 | 813 | $402 \pm 16$ | 0.81 |
| cefoxitin | 757 | 608 | $304 \pm 15$ | 0.8 |
| nitrofurantoin | 713 | 559 | $287 \pm 12$ | 0.78 |
| cefotetan | 124 | 95 | $49 \pm 6$ | 0.77 |

The specific proteins transferred between chromosomes are known to vary by ICE feature. This is demonstrated in part by the AMR proteins analyzed in Figures 5 and 6. It is possible to classify these features in a general way considering all ICE features used in this study. Figure Genome-IPR Co-occurrence Map shows the cooccurrence of ICE proteins by genome, for the 6000 genomes with the highest cargo protein fraction. To this set, 500 genomes containing the largest number of rare ICE features were added to ensure representation of those families defined by a smaller collection of well-characterized proteins from the literature (see Table 4. Figure 7 thus represents the co-occurrence of ICE features within the selected set of genomes. From the co-occurrence matrix, one can calculate a distance between all *pairs of genomes* using the Euclidean distance between their representation as normalized ICE feature vectors. The resulting genome-genome distance map, shown in Figure 8, provides a hierarchical clustering of genomes based on the co-occurrence of ICE features. In principle, one can use the same vectorization procedure on genomic properties other than ICE features, defining a different set of genomes of interest within the *mobilome*, to (re)classify organisms not by name, but by distance in a space of mobilization features. Note that only 1 in 75 genome labels (by genera) are shown along the y-axis given the relatively large number of genomes included.

**Figure 8.**
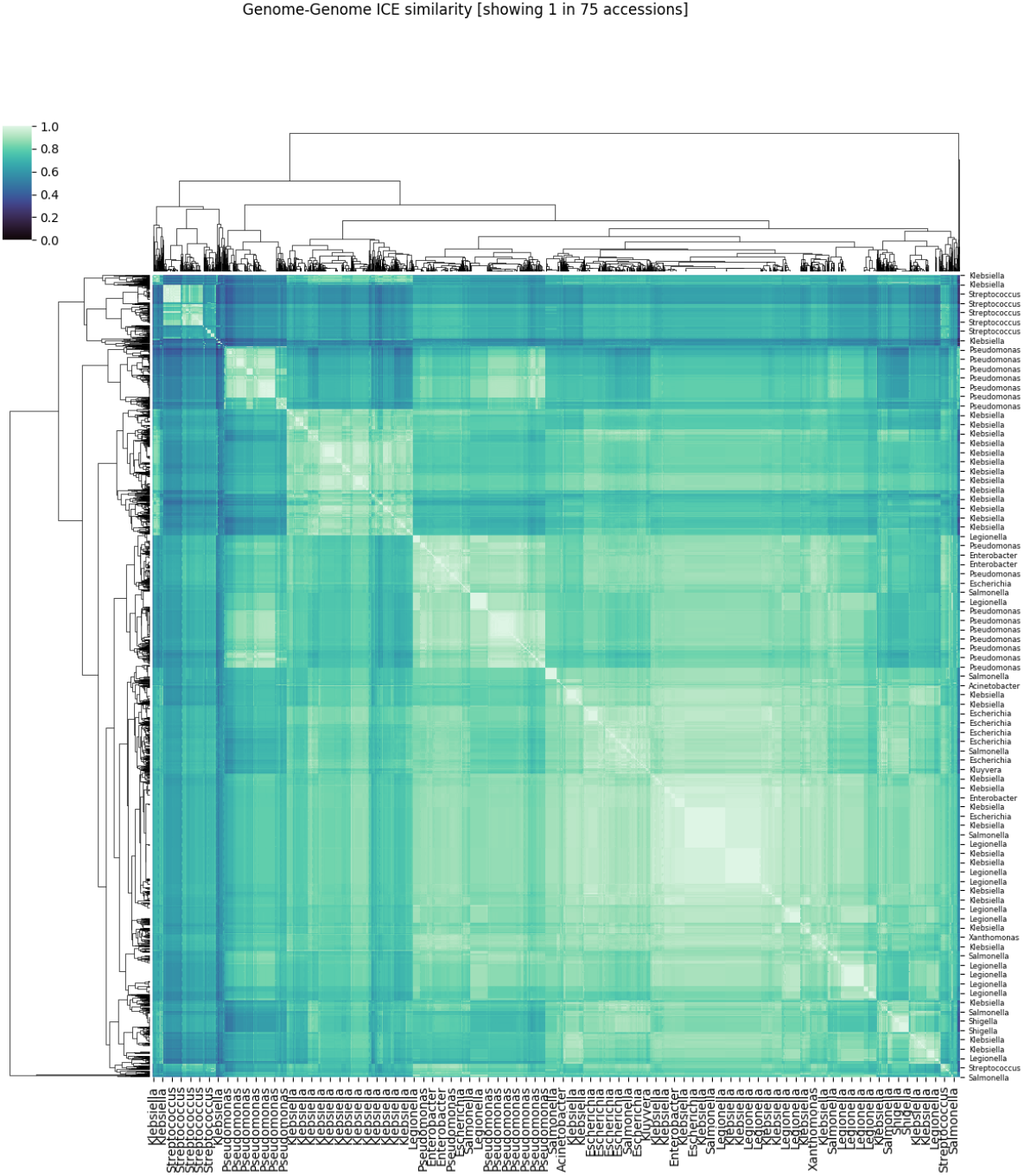
Heatmap of Genome-Genome Distances. Genome-Genome distance based on Euclidean distance between vectors of ICE features for the 6500 genomes used in 7. One in 75 labels are rendered on each axis. The figure shows how different sets of genomes cluster based on different co-occurring ICE features (see text).

**Figure 7.**
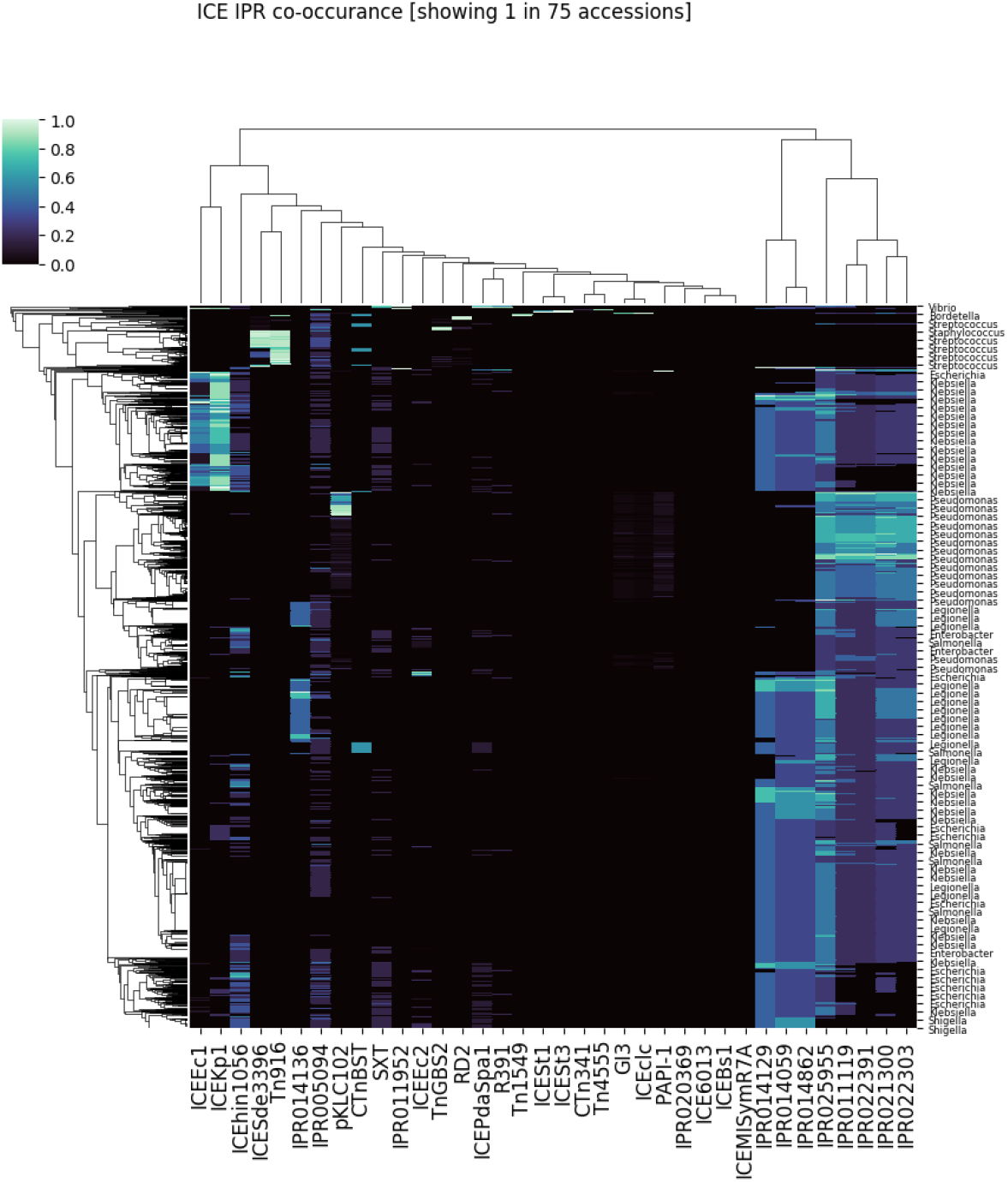
Genome-IPR Co-Occurance Map. The co-occurrence of ICE features by genome, for the 6000 genomes with the highest cargo protein fraction and 500 genomes containing rare ICE features (see text). Co-occurrence of particular ICE features does vary with taxonomic group.

**Table 4.**
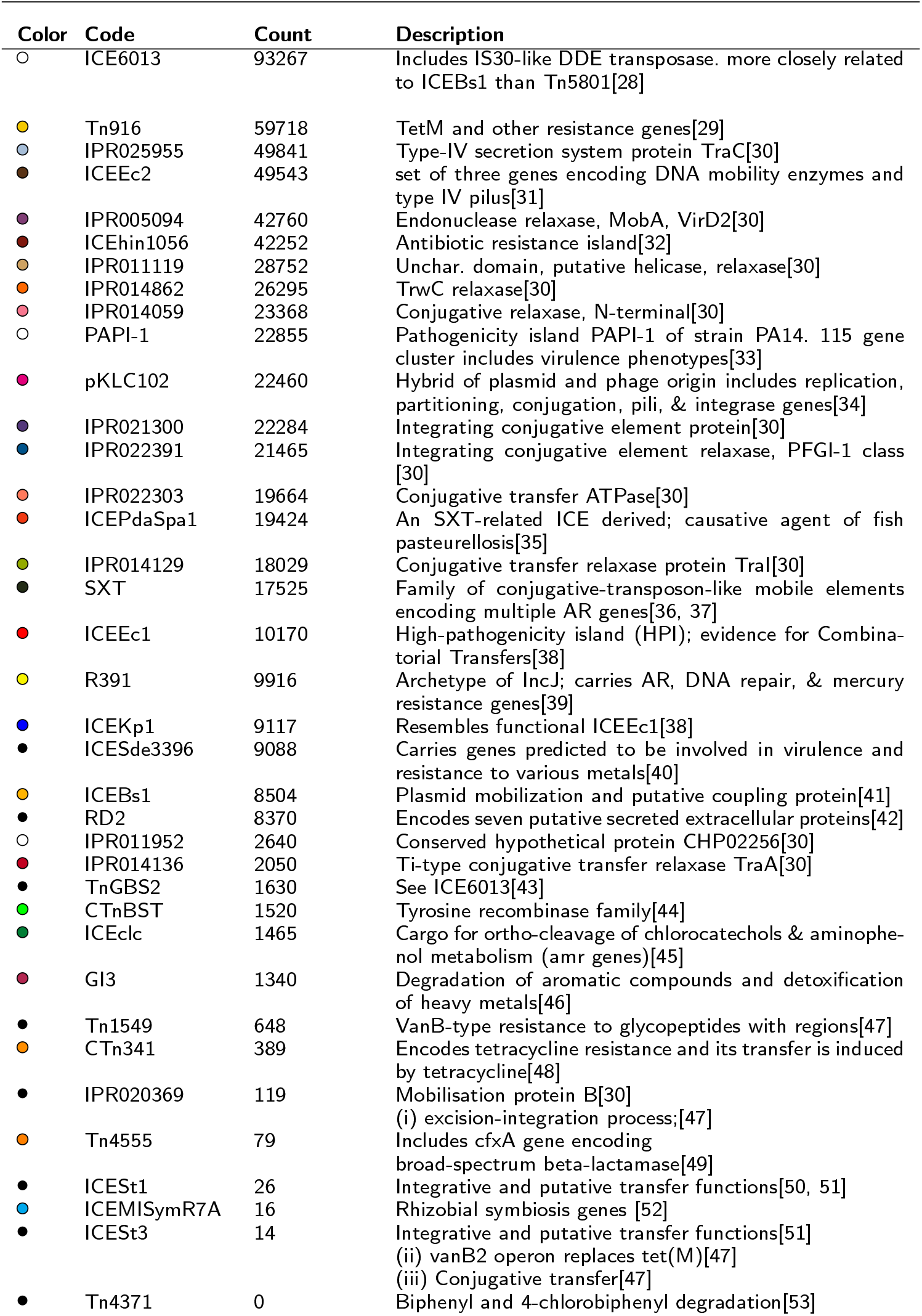
Table of ICE features included in this study. The first column indicates the pixel color representing ICE features observed in Figure 1. The genome count represents the number of genomes that contain the corresponding ICE feature.

## Discussion

The public availability of large scale genomic data makes it possible to apply cloud computing technology and big data techniques to the study of important phenomena in molecular and microbiology. Furthermore, putting all of this data in a relational database with biologically structured entity relations (i.e., linking genomes, genes, proteins, domains, and metadata) provides a powerful new way to ask biological questions about the data. We leveraged this approach in the current study of ICE and associated cargo proteins. Exchange of proteins by conjugative processes, including ICE, is now understood to be an essential mechanism by which bacteria acquire new phenotypes, transmit molecular functions, and adapt to stress. Furthermore, these events are critical for understanding bacterial evolution and phylogeny [15, 11, 12]. Our work not only sheds light on ICE-cargo protein transfers between and within genera, but also demonstrates the power of a “big data” approach for improving our understanding of bacterial evolutionary dynamics, particularly surrounding important phenotypes such as AMR.

In our analysis we identified sets of proteins with the strongest evidence as ICE and ICE cargo proteins. This was accomplished by selecting only those proteins that exhibited both 100% sequence identity and co-occurrence *in pairs of genomes* containing identical ICE sequences. With this strict selection process, the putative cargo proteins exhibited a high degree of spatial correlation within assembled contigs (i.e., they were highly adjacent to each other, as well as *to the ICE protein itself*). Other proteins in these genomes may also have been transferred (or are transferable) by ICE, but they did not meet our strict selection criteria. Considering only these candidate cargo proteins, we were able to profile the frequency of ICE-mediated protein exchange within and between genera.

Our results suggest that ICE-mediated exchange is not uncommon [12, 11]. ICE proteins are observable in 51% of bacterial genomes and in 626 of 1,345 genera (47%). Rates of intra- and inter-genus ICE-mediated exchange varied significantly depending on the taxa involved, suggesting that taxonomy is a significant “risk factor” for genetic exchange of, e.g., AMR or pathogenicity proteins [22]. By quantifying this risk across a large database of high-quality WGS data, we measured the “exchange likelihood” between different taxa, and visualized these frequencies as a force-directed graph (Figure 3). Our analysis revealed distinct clusters of genera with high rates of cross-genus ICE-mediated exchange. This suggests that the likelihood of protein transfer varies substantially by genus pair, and that the bacterial composition within a given environment is an important consideration when attempting to evaluate mobilization potential within a microbial community. In practical terms, this means that analysis of the risk posed by the commensal microbiome (i.e., as a potential reservoir of AMR) must take into account the specific composition of differing microbiomes, including the presence of bacterial taxa that are more likely to engage in genus-genus exchange with pathogens of interest. In other words, there is no “one-size-fits-all” mobilization metric [14]. While we have conducted this analysis for ICE features, the analytic approach can be applied to any mobile genetic element(s) and cargo protein(s) of interest. In this way, our overall approach represents a method for obtaining a long-range evolutionary view of transfer likelihood between diverse bacterial taxa, including pathogens and commensal bacteria [11]. These “baseline” exchange likelihoods are critical parameters for risk analysis at the microbial community level [13, 23].

Bacterial taxon is not the only significant driver of exchange likelihood; we have also observed that putatively successful transfer events are more likely to involve cargo proteins that infer fitness advantage to the involved bacterial populations, such as AMR. While any gene can, in principle, be transmitted as a cargo gene in conjugative exchange, only a subset of transferred proteins will increase the fitness of the receiving organism. The likelihood of observing successful transfer depends on a large number of factors including the environment, the existing proteins in the recipient chromosome, the cargo proteins themselves, and the survival probability of the organism [24].

ICE protein transfer that improves fitness may increase survival probability. Therefore, chromosomal arrangements that group *fitness-conferring* cargo proteins near the ICE machinery will be observed more frequently than those arrangements that involve neutral or disadvantageous proteins. Conversely, very common proteins that aid in stress response may be less likely to be transferred as cargo, since the relative fitness advantage is diminished for proteins that are already likely to be present within a bacterium. The particular stressor – as well as the specific advantageous stress response proteins of interest – depend on phenotype of interest. This view is exemplified by the data in Table 2 which shows that *rare* AMR proteins are more likely to be found as cargo in ICE versus non-ICE genomes. Conversely, common AMR proteins are less likely to be found in ICE genomes. One might hypothesize that, with chromosomal rearrangement, nature effects a real world *“Monte Carlo”* experiment to dynamically optimize cargo protein collections - thereby spreading rare (but useful) proteins and gene combinations over time.

The particular proteins transferred between chromosomes is known to vary with the different features of ICE, as demonstrated by the AMR proteins analyzed in Figures 5 and 6. Considering all ICE features used in this study, Figure 7 shows the co-occurrence of ICE proteins by genome for the 6000 genomes with the highest cargo protein fraction. To this set, 500 genomes containing the largest number of rare ICE features were added to ensure representation of those families defined by a smaller collection of well characterized proteins from the literature (see Table 4. Each row of this matrix corresponds to one genome, labeled not by accession but by genus. These results suggest that ICE dynamics are important in structuring genomic content, and thus driving phylogenetic evolution. Based on Figure 7, it seems that sometimes these evolutionary ICE dynamics overpower other taxonomic drivers, such that genus-level genomes do not cluster together. To demonstrate this ICE-driven phylogeny, we used the data in Figure 7 to generate Figure 8, which represents the distance between all pairs of genomes based on Euclidean distance between their representations as normalized ICE feature vectors. The resulting hierarchical clustering shows that the dominant ICE features include genomes across different genera and, conversely, that individual genera include genomes representative of different ICE features. This abrogation of genus-level taxonomy due to ICE-related genomic content is an inevitable consequence of the cross-genus transfers visualized in Figure 3 and the corresponding Force-Directed graph of ICE transfer. Given the reality of conjugative exchange, there is no reason to expect that taxonomic classification by organism name will always predict the composition of ICE cargo. However, by selecting genomes based on a particular phenotype of interest, it is possible to classify organisms and genome-genome distances based on a feature space defined by ICE (or other mobilization) proteins. Given the ubiquity and diversity of ICE and other types of conjugative exchange [11], these types of genome clustering techniques may provide crucial information about bacterial evolution that is not contained within traditional phylogenies.

This approach may also be useful in exploring the role of HGT in bacterial evolution as it related to AMR mechanisms in particular. The environments in which bacteria live, the stresses they encounter, and their genetic composition are all dynamic. If the proteins required for successful response to a commonly encountered environmental stress are themselves common, then the fitness advantage gained by maintaining those proteins as ICE cargo is diminished. If the environmental stress is relatively new (e.g., a new antibiotic compound), and the protein(s) required for survival against this antibiotic are rare, then the right combination of proteins may significantly increase the organism’s survival probability and, therefore, the likelihood of transmitting (and of observing) those proteins as ICE-associated cargo is also increased. Under this hypothesis, the ICE process is an important mechanism for spreading *new or less common* stress responses and resistance mechanisms; but is relatively unimportant for maintaining the genetic material required for stress response in common or oft-encountered environments. Given that the widespread use of most antibiotic compounds is a relatively new phenomenon (at least in evolutionary terms), this may explain why AMR genes that are specific to antibiotic drug compounds are over-represented within ICE genomes, while more generalist AMR mechanisms are over-represented within non-ICE genomes. Furthermore, this may explain why the resistance phenotype for all analyzed antibiotic drugs was much more likely to be associated with isolates containing ICE feature(s), compared to isolates not containing ICE features.

## Conclusions

By using a big data approach on both genotypic and phenotypic NCBI data, we generate important insights into bacterial population-level genetic exchange and evolution, and demonstrate the importance of microbial composition in the likelihood of ICE-mediated transfer events. Because our analysis is based on over 166,000 curated and high-quality WGS datasets from NCBI, it can serve as the basis for understanding differences in the ICE transfer likelihood of specific cargo genes between specific bacterial genera. These “baseline likelihoods” are crucial to quantifying the likelihood of pathogens obtaining genetic material (e.g., AMR genes) from commensal microbes residing in the same environment. Furthermore, we demonstrate that the importance of ICE-mediated exchange may differ based on the relative rarity of the cargo gene, suggesting that ICE-containing genomes may be an important target for surveillance of emerging phenotypes. Given the human and public health importance of AMR, virulence and pathogenicity, our results therefore provide an important foundation for improved quantitative assessment of the microbial risk posed by various microbiomes.

## Methods

### Data Description

#### NCBI Sequence Data

NCBI maintains a large, public domain repository of raw WGS data. As described in *Genome Curation and Selection*, we retained 186,887 (non-redundant) public genomes, which included raw Illumina paired-end bacterial sequence data from the Sequence Read Archive (SRA), and high quality assembled genomes maintained in the RefSeq Complete genome collection, genbank, and NCBI’s pathogen tracker. A detailed description of the genome curation, assembly, and annotation pipelines may be found in methods, the online supplement, and in a recent paper by Seabolt et al. [25, 26, 27]. This selection provided us with genomes representing 1,345 genera of bacteria. Accession identifiers for all of these data sets are available in the supplement. In order to obtain evidence for candidate cargo proteins (i.e., proteins that had potentially been transferred between bacteria via ICE-mediated HGT), it was necessary to first identify genomes containing known ICE proteins. We further restricted this set by considering only sets of genomes that shared particular ICE proteins with identical amino acid sequence (hereafter referred to as *ICE genomes*). For those specific genomes sharing identical ICE proteins we then searched for *other* proteins (with sequence identity). These are then labeled possible *cargo*.

This calculation is of order *O*(*N*^2^), but is straightforward to compute on any large cluster or cloud infrastructure. However, due to the high compute cost, we chose to first put the required intermediate artifacts into a relational database we call OMXWare [27]. With an appropriate schema, the database links each genome to all of its relevant unique proteins, domains, and functional annotations, as described in *Genome Assembly and Annotation*. Analysis of ICE genomes and candidate cargo proteins then simply becomes a set of database queries. This process not only reduces final storage requirements (since the unique gene and protein tables are non-redundant), but also makes the large-scale computation and database reusable for future biological studies.

#### NCBI Antibiotic Susceptibility Testing BioSample Data

To analyze associations between phenotypic AMR and ICE, we retrieved metadata for each NCBI accession that contained antimicrobial susceptibility testing (AST) data. Retrieved metadata included genomic accession number for each isolate, as well as the antibiotic compound against which it was tested, the test type and the phenotypic outcome (resistant, susceptible, or intermediate). We considered only those isolates with a “resistant” phenotypic outcome to be resistant. By linking the BioSample accession with the SRA accessions in OMXWare, we were able to identify genomes for which corresponding AST data are available. These genomes were used in our analysis of phenotyic AMR and ICE.

### Genome Assembly and Annotation

All of the bioinformatic tools used are open source. To assemble Whole Genome Sequence from the SRA, Trimmomatic 0.36 [54] is used to remove poor-quality base calls, poor-quality reads and adapters from the sequence files. For removal of PhiX control reads, Bowtie 2 2.3.4.2 [55] is used to align the sequences to references derived from PhiX174 (Enterobacteria phage phiX174 sensu lato complete genome). FLASh 1.2.11 [56] is used to merge paired-end reads from the sequences to improve quality of the resulting assembly. Once these pre-assembly steps are complete, the files are passed through SPAdes 3.12.0 [57] and QUAST 5.0.0 [58] in an iterative assembly/quality evaluation process. After assembly, genes and proteins are annotated using Prokka 1.12. [59] Following gene and protein annotation, protein domains are determined using InterProScan 5.28-67.0 [60]. All 16 available analyses provided by InterProScan are run over all input sequences. Results are output in JSON format. For each of the 16 resulting JSON documents produced by InterProScan, we parse the annotated domain information into a set of delimited files which are then loaded into the appropriate structured tables in a DB2 database.

Due to the large scale of sequence data in this study, we implemented a cloud-based architecture to effectively orchestrate the complex use of the bioinformatic tools described above across multiple servers. These tools and the detailed architecture are described in the supplement. The bioinformatic tools were provisioned and deployed on the IBM Cloud utilizing a combination of bare metal and virtual machines totaling over 1468 CPUs, 6TB RAM, and 160TB of hard drive space. IBM Spectrum Scale version 5 was used for cluster filesystem access. Apache Mesos version 1.6.1 provided cluster resource management and scheduling. Marathon version 1.6.352 was used for container orchestration and health checking with all deployed processes running as Docker containers. The system utilizes a message-oriented methodology for executing pipelines using RabbitMQ 3.7.2 as a broker. System events emitted from all pipelines are captured in the timeseries database InfluxDB 1.2.0, and annotated information was stored in DB2 v11.1.3.3. A core component in our architecture is the OMXWare Distributed Pipeline Framework (ODPF). This component coordinates incoming messages from the broker, executes individual stages of a pipeline, records events as each stage of a pipeline progresses, and routes messages to additional queues when requested. The supplement provides additional details about the architecture including pipeline execution mechanics and annotation processing. This annotation process identified a total of 66,945,714 unique gene sequences and 51,362,178 unique protein sequences, yielding 138,327,556 unique protein domains along with related functional annotations (e.g., IPR codes).

### Genome Curation and Selection

OMXWare systematically curates WGS files from NCBI because the accuracy of metadata and quality of WGS files maintained by NCBI varies dramatically. Some files, for example, may have been derived from contaminated samples and are therefore not technically WGS files, while others may be labeled improperly with the incorrect microbial genus ID.

Bacterial SRA datasets that were at least 100 MB of size were downloaded and converted to FASTQ format using SRA-tools [25]. Genome assemblies containing greater than 150 contigs (of size > 500 bp) and an N50 of less than 100k bp were discarded with the exception of genomes from the genus Shigella, where assemblies containing greater than 500 contigs (of size > 500 bp) and an N50 of less than 15,000 bp were discarded. From the original assemblies (at the time of this study), 186,888 genomes passed the quality criteria described above. A detailed description of the entire assembly, annotation, and InterPorScan process is reported elsewhere [27].

### ICE and Cargo Protein Selection

ICE proteins are still being discovered. Large families of such proteins are known in the literature [61, 3, 4] and those used in this study are enumerated in Table 4. In addition to these features, InterProScan implements a coding system for classification of proteins [60]. The complete set of Interpro codes used in this study is listed in Table 4. These codes describe conserved domains or larger families of proteins that provide essential function in the ICE transfer process including conjugative relaxases, nickase, helicase, and other mobilisation proteins.[62, 63, 64, 65] Relaxases are DNA strand transferases that bind to the origin of transfer (oriT) in the ICE transfer process and melt the double helix.[66, 64] Helicase is required to separate double-stranded DNA into separate strands for duplication. Nickases introduce single-strand breaks in DNA. Together, these proteins define a *relaxation complex* or *relaxosome*. In some cases, several different domains on the same protein will each contribute distinct function required for ICE. These InterPro (IPR) codes assigned to ICE related proteins are also listed in Table 4. [67, 61, 60]. These groups of integrative conjugative element proteins exhibit significant sequence diversity [1, 2, 5, 6, 7], but represent conserved domains involved in the machinery required for ICE. Of the 186,888 bacterial genomes processed, 95,781 genomes (~51%) were found to contain a protein belonging to one or more of the ICE families listed in Table 4. The genomes with one or more ICE feature contained 405,065,204 total proteins, representing 26,491,056 unique protein sequences.

There are a total of twelve Interpro codes associated with ICE listed in Table 4. The IPR005094 family (exemplified by MobA/VirD2) describes relaxases and mobilisation proteins. [62, 63, 60] The code IPR011119 represents a domain found in a family of proteins in proteobacteria annotated as helicase, conjugative relaxase or nickase.[68, 60] IPR014059 codes for a domain in the N-terminal region of a relaxase-helicase (TrwC) that acts in plasmid R388 conjugation. It has been associated with both DNA cleavage and strand transfer activities. Members of this family are frequently are *“near other proteins characteristic of conjugative plasmids and appear to identify integrated plasmids when found in bacterial chromosomes*”.[64, 60] The IPR014129 family represents proteins in the relaxosome complex (exemplified by TraI). TraI mediates the single-strand nicking and ATP-dependent unwinding of the plasmid molecule. The two activities are driven by separate domains in the protein. [66, 60] IPR014862 represent a conserved domain found in proteins in the relaxosome complex (exemplified by TrwC). [64, 60] The IPR021300 family represents a conserved domain observed in ICE elements in the protein family PFL_4695 (originally identified in Pseudomonas fluorescens Pf-5). [61, 60] IPR022303 describes a family of conjugative transfer ATPase representing predicted ATP-binding proteins associated with DNA conjugal transfer. They are found both in plasmids and in bacterial chromosomal regions that appear to derive from integrative elements such as conjugative transposons (so they may not be unique to ICE). IPR025955 describes a family of TraC-related proteins observed in Proteobacteria. TraC is a cytoplasmic, membrane protein encoded by the F transfer region of the conjugative plasmid. It is also required for the assembly of the F pilus structure. The family includes predicted ATPases associated with DNA conjugal transfer. [65, 60] IPR022391 represents a conjugative relaxase domain in the PFGI-1 class. Proteins with this domain include TraI putative relaxases required for ICE and found in Pseudomonas fluorescens Pf-5. They are similar in function to TraI relaxases of the F plasmid, but have no sequence homology. This Interpro entry represents a N-terminal domain of proteins in this class.[60] IPR025955 represents a family of conserved Type-IV secretion system proteins, TraC/Conjugative transfer ATPase, in Proteobacteria. TraC is a encoded by the F transfer region of the conjugative plasmid and is required for construction of the F pilus structure. The filamentous F pili are serve to create and maintain physical contact between conjugating donor and recipient cells. The family also includes predicted ATPases associated with DNA conjugal transfer. They are found in ICE elements. [65, 60]

Taking advantage of the relational database, which contains tables that associate protein UIDs with INTERPRO ACCESSIONs, we used SQL queries first to obtain a full list of UIDs for all the proteins which matched one or more of the features described above and listed in table 4. This exhaustive list produced a list of 28,042 *candidate ICE proteins*, from which we then removed all those that appear in only 1 genome (by sequence identity). As part of this culling step, performed using SQL and which resulted in 15,398 *ICE proteins* associated to the selected IPR codes or features, and appear in at least two genomes (see Table 5), we persisted the containing 95,781 genomes, which we call *ICE genomes*. To determine the percentage of ICE proteins in these ICE genomes, we also obtained counts of all distinct proteins that these genomes contain, 21,207,794.

**Table 5.** ICE-Related Statistical Counts

| Description | Unique Sequences |
| --- | --- |
| Total genomes examined | 186,887 genomes |
| Genomes containing an ICE protein | 95,781 genomes |
| Candidate ICE proteins | 28,042 amino acid |
| All proteins in ICE genomes | 21,207,794 amino acid |
| ICE proteins found in 2 or more genomes | 15,398 amino acid |
| Distinct cargo proteins in 2 or more ICE genomes | 11,276,651 amino acid |
| Cargo transfer count within ICE genomes | 4,938,737,476 amino acid |

Upon identification of the ICE proteins and corresponding containing genomes, further SQL queries were used to identify proteins most likely to be cargo proteins based on evidence of ICE transfer. To make this selection we queried for proteins that are not only present in the same genome pair as ICE proteins, but with the added restriction that they not be present in any non-ICE genome. For this purpose we first queried the database for all proteins in the 95,781 ICE genomes, which returned 387,682,038 distinct <ICE genome accession number, protein UID> tuples. In many cases a unique sequence is observed in more than one genome; in total there were 21,207,794 distinct protein sequences in the set of ICE genomes. This group was then filtered to identify the subset of distinct proteins appearing in two or more (ICE) genomes. If a protein sequence is seen only once, then by definition there is no supporting evidence of it being “transferred”. To further reduce false positive identification of transfer by ICE (vs being vertically transferred), we discard any protein that appears in any of the 99,052 *non-ICE genomes*, thus effectively establishing a rigid approach whereby proteins of interest must exclusively appear in ICE genomes. With this strict selection process, we identified 11,276,651 distinct sequences that are associated with ICE, have evidence of transfer, and lack evidence of transfer in a non-ICE genome. We refer to this set as *cargo proteins*, i.e. proteins with the greatest evidence of transfer in an ICE process. Furthermore, tabulating the number of <cargo protein, ICE genome A, ICE genome B> triples where ICE genome A and ICE genome B contain at least one identical ICE protein sequence, yields a total of 4,938,737,476 transfers. The data is summarized in Table 5

### Estimating False Positives: Classification of Plasmid Proteins

Some of the features used to identify families of proteins indicative of ICE correspond to proteins that mediate required for Integrative Conjugative Exchange. However, some may also be required for the formation (or integration) of plasmids. In order to measure or bound the possible rate of false ICE classification, all bacterial plasmids were downloaded from NCBI, and the annotated proteins placed in a database table of plasmid proteins. The MD5 Hash was used as the primary key for entries in this table. The MD5 hash is used as the primary key for every sequence entity in the database. Determining if a protein has been observed on a plasmid in the NCBI reference is then accomplished by querying if the primary key of the protein in question exists in the table of plasmid proteins. This data was used in the analyses shown in Figure 4.

### Characterizing Cargo Proteins

Once all candidate ICE and cargo proteins were identified, the database was then queried to obtain annotated protein names, along with the genomes and genera in which they were observed. This data was saved to a delimited file, which was then used as input to python scripts to obtain a list and count of intra- and inter-genera cargo protein transfers, per protein name. This O(n2) algorithm required an exhaustive pairwise comparison of genomes, detecting initially the intersection of ICE proteins between each genome pair and, if non-empty, the pair’s common cargo proteins. This process initially identified a total of 5956 ICE-related genus-genus transfers (i.e. triples of the form (genus1, genus2, protein-name), carrying a total of 23,353,196,048 exchanged protein sequences. The resulting count is non-distinct by protein name as, for instance, Tyrosine recombinase XerD is transferred both between Salmonella-Salmonella genomes, as well as between Oligella-Proteus genomes and other genera pairs. Of these triples, 1680 transfers are observed to occur between genome pairs belonging to the same genus, carrying a total of 23,242,566,032 sequences. Additionally, counts of per protein transfers are similarly maintained, by genus. Note that this approach does not allow us to determine transfer direction, therefore no determination was made regarding source and target.

### Intra and Inter-genus protein transport

Inter-genus protein transport was represented as a graph in which each genus is a node, and an edge between a pair of nodes represent a value of co-occurrence of <ICE protein, cargo protein> pairs, as a Force Directed Graph shown in Figure 3, discussed in *Results*. The Force Directed Graph is also available as a simple Web Application that allows users to select which ICE families and or types of Cargo genes to display.

### ICE and Cargo Proximity

To test the loci proximity of ICE and cargo genes in the discovered genome pairs, we utilized our compiled list of genera, genomes, ice and cargo proteins, and leveraged Prokka’s accession index (a positional indicator of a gene or protein’s sequence in a particular genome). While this approach is limited by the fact that assembly-based contigs are unordered, and therefore introduce gaps in the genome sequences in our database, data from the accession gene or protein index indicates sequential placement of ICE and cargo proteins (Figure 1).

### Characterizing AMR Genes

Once all candidate ICE and cargo proteins were identified, the database was then queried to obtain annotated protein names, along with the genomes and genera in which they were observed. To gain insight into putative cargo that co-occurs with ICE proteins but not yet annotated as ICE, we looked in particular at the cargo proteins with AMR names. To accomplish this we used the MEGARes database, which contains AMR gene sequences with associated unique identifiers, as well as a hierarchical ontology to classify each AMR gene [18]. All unique sequences in MEGARes were run through the same pipeline used to annotated the set of all proteins in OMXWare. This provides a self consistent annotation. We then obtained all putative cargo proteins with names that matched any AMR protein name. This data was used to compute, for every AMR name, the frequency of observation as possible cargo in an ICE genome and the frequency of observation in genomes not associated with ICE.

### Tabulation of Data

To perform key analysis reported here, data was first tabulated in database tables or views (and exported to delimited files for the supplement). ICE genomes (defined above) were tabulated by Genus as shown in Table 1. Observed ICE features from Table 4 were tabulated by Genome for the analysis shown in Figures 2 and 7. Similarly, proteins assigned names associated with antimicrobial resistance were tabulated by number of ICE and non-ICE genomes, and the fraction of unique sequences observed in plasmids were tabulated as well (See Figure 2).

### Hierarchical Clustering and Co-occurrence

The proteins studied here, and the ICE families themselves, are not in-dependant features. Hierarchical clustering was used to characterize the correlations an cooccurrence of proteins or genomes by ICE family (see Figures 7, 5, and 6. This was accomplished by forming a co-occurrence matrix and using the Seaborne *clustermap* algorithm which does single linkage clustering to form heatmap and dendrogram relating the co-occurring features [69]. The vector is ICE features for an entity (protein or genome) are then be used as a Euclidean metric (dimensionality reduction) to compute an effective distance between the entities. This approach was used to cluster and visualize the genome-genome distance shown in Figure 8. Seaborne *clustermap* was used for this step as well.

## Author’s contributions

J.H.K. and I.T. performed and designed research, compiled supporting data, analyzed data, and wrote the paper. G.N. and A.A. developed big data visualizations, classified AMR genes and assay data, and contributed to the paper. E.S. and M.R. built the database, pipeline, and contributed to the paper. N.N. proposed the study, designed the research, and wrote the paper.

## Acknowledgements

The authors would like to acknowledge contributions from M. Roth, H. Krishnareddy, K. Clarkson, and E. Kandogan.

